# Age differences in neural distinctiveness during memory retrieval versus reinstatement

**DOI:** 10.1101/2023.03.21.533591

**Authors:** Claire Pauley, Malte Kobelt, Markus Werkle-Bergner, Myriam C. Sander

## Abstract

Robust evidence points to mnemonic deficits in older adults related to dedifferentiated, i.e., less distinct, neural responses during memory encoding. However, less is known about retrieval-related dedifferentiation and its role in age-related memory decline. In this study, younger and older adults were scanned both while incidentally learning face and house stimuli and while completing a surprise recognition memory test. Using pattern similarity searchlight analyses, we looked for indicators of neural dedifferentiation during retrieval and asked whether this might explain interindividual differences in memory performance. Our findings revealed age-related reductions in neural distinctiveness during memory retrieval as well as in encoding-retrieval reinstatement in visual processing regions. We further demonstrated that the degree to which patterns elicited during encoding were reinstated during retrieval tracked variability in memory performance better than retrieval-related distinctiveness only. All in all, we contribute to meager existing evidence for age-related neural dedifferentiation during memory retrieval. We propose that the recognition task (as opposed to a cued recall task) may have revealed impairment in perceptual processing in older adults, leading to particularly widespread age differences in neural distinctiveness. We additionally provide support for the idea that well-defined reactivation of encoding patterns plays a major role in successful memory retrieval.

## 1 Introduction

Cognitive decline in neurologically healthy aging is frequently associated with a phenomenon called age-related neural dedifferentiation (for reviews, see Koen & Rugg, 2019; Koen et al., 2020; Sommer & Sander, 2022). Findings of neural dedifferentiation have been interpreted in relation to neurobiological models suggesting that cognitive aging results from impaired neurotransmitter function, predominantly affecting the dopaminergic system (S-C Li et al., 2001; S-C Li & Rieckmann, 2014). These models propose that as a result of insufficient dopamine modulation, neural information processing in older adults suffers from poor signal transmission leading to increased neural noise and reduced neural distinctiveness. In functional magnetic resonance imaging (fMRI) studies, neural distinctiveness is often operationalized by contrasting content-related neural activity between different visual categories (e.g., faces and houses) either using mean blood-oxygen-level-dependent (BOLD) signal (e.g., Park et al., 2004) or using multivoxel activity patterns (e.g., Hill et al., 2021). These studies have accumulated evidence for age differences in the distinctiveness of neural signals (S-C Li et al., 2001; DC Park et al., 2004; for review, see Koen & Rugg, 2019).

Recently, the impact of age-related neural dedifferentiation on episodic memory performance has received particular interest due to the significance of highly specific neural representations for encoding and retrieving distinct events. Several studies have established a relationship between low representational specificity during *memory encoding* and memory decline in older adults (Zheng et al., 2018; Koen et al, 2019; Srokova et al., 2020), suggesting that the formation of well-defined, non-overlapping memory traces is important for memory performance. Most evidence examining neural dedifferentiation during *memory retrieval* comes from assessments of age differences in cortical reinstatement. The *cortical reinstatement hypothesis* suggests that cortical representations of information during memory retrieval are reactivated mirror-images of their respective representations during encoding (Norman & O’Reilly, 2003; Johnson & Rugg, 2007; for review, see Danker & Anderson, 2010). Age-related declines in the precision of cortical reinstatement have been reported at both the item (St-Laurent et al., 2014; Bowman et al., 2019; Folville et al., 2020; Hill et al., 2021) and category (McDonough et al., 2014; Johnson et al., 2015; Abdulrahman et al., 2017; Bowman et al., 2019; Deng et al., 2021; Hill et al., 2021) representational levels (but, see Wang et al., 2016; Thakral et al., 2017; Thakral et al., 2019, for absent age effects, and Deng et al., 2021, for age-related hyperdifferentiation). These age differences in reactivation have frequently been associated with senescent memory decline (St-Laurent et al., 2014; Abdulrahman et al., 2017; Bowman et al., 2019; Hill et al., 2021) underlining the mnemonic advantage afforded by strong representational distinctiveness in cortical reinstatement.

Recent evidence has suggested that, in addition to cortical reinstatement, neural representations during memory retrieval may also reflect spatial transformations of the representations initially formed during encoding (Xiao et al., 2017; Favila et al., 2018; for review, see Favila et al., 2020). For example, Xiao and colleagues (2017) demonstrated that the representational structure of stimuli in the ventral visual cortex during encoding were reactivated in the frontoparietal cortex during retrieval, indicating a spatial shift in content-related representations between encoding and retrieval. Furthermore, the integrity of the representational structure was best maintained through this transformation in comparison to reactivation within the ventral visual cortex. These findings suggest that neural representations supporting memory retrieval may not solely mirror encoding processes, but rather flexibly adapt to serve retrieval demands. This idea was supported by Favila and colleagues (2018) who manipulated the retrieval goal, such that they explicitly asked participants to retrieve the stimulus color and the stimulus object in separate retrieval trials. They found that both features were reliably reinstated in the occipitotemporal cortex regardless of retrieval goals, but the lateral parietal cortex represented the stimulus color during color trials and the stimulus object during object trials. In other words, activity patterns in the lateral parietal cortex representing retrieved content flexibly adapted to facilitate the current retrieval goal. This raises the question of whether manipulations of task goals or attention during retrieval may reveal additional regions of distinctiveness susceptible to age-related decline in addition to those observed during encoding-related reinstatement. So far, the literature on age-related neural dedifferentiation has largely ignored this possibility, which has focused almost exclusively on the reduced specificity of cortical reinstatement in older adults. A notable exception comes from Hill and colleagues (2021) who demonstrated that older adults also reveal deficits in category-level distinctiveness during retrieval, not only during encoding. However, the analysis was restricted to two pre-defined regions of interest. As the transformation literature suggests (for review, see Favila et al., 2020), content-related representations during retrieval may lie outside the bounds of encoding representations. Accordingly, the investigation of potential age-related impairment during retrieval requires methods that allow for the exploration of neural patterns across the whole brain. To date, few studies have considered the impact of age-related neural dedifferentiation on memory retrieval independently of reinstatement effects using a whole-brain approach (Dulas & Duarte, 2012; St-Laurent et al., 2014; Johnson et al., 2015). All studies found evidence for an age-related decline in neural distinctiveness during retrieval. Crucially, both St-Laurent and colleagues (2014) and Johnson and colleagues (2015) reported age differences in neural distinctiveness during retrieval that could not be attributed to age differences during encoding, indicating that dedifferentiation during retrieval may not be fully captured by only looking at reinstatement.

There is substantial evidence suggesting that distinctive cortical reinstatement benefits memory performance (St-Laurent et al., 2014; Abdulrahman et al., 2017; Bowman et al., 2019; Hill et al., 2021). However, it is unclear whether the distinctiveness of representations during retrieval supports memory performance solely through reinstatement of encoding content or perhaps serves as an additional boost. In the aging brain, it has been suggested that less precise neural signaling during retrieval may have a compounded negative effect on mnemonic content already suffering from reduced distinctiveness during encoding (Sander et al., 2021). Accordingly, memory deficits in older adults may be even better explained by age differences in neural distinctiveness during retrieval than during encoding or reinstatement. Few studies have examined the link between reduced neural distinctiveness during retrieval and memory performance. Dulas and Duarte (2012) did not identify any brain regions demonstrating memory-related effects associated with age-related neural dedifferentiation for objects or words during memory retrieval. However, their search for a voxel-wise main effect using a univariate approach may not have been sensitive enough to pick up on mnemonic content stored in fine-grained neural activity patterns. Using a multivariate decoding model, Johnson and colleagues (2015) successfully linked distinctive retrieval representations to subjective vividness ratings in both younger and older adults. Their findings showed that distinctiveness in the prefrontal cortex was more closely associated with subjective vividness in older adults compared with younger adults and that distinctiveness in the parietal cortex tracked subjective vividness better in younger adults compared with older adults. Subjective assessments of memory often differ from objective findings (Johnson, 2006), especially in older adults (Norman & Schacter, 1997). It therefore remains an open question as to whether retrieval-related neural distinctiveness reflected in multivariate representations would demonstrate a relationship with an objective measure of memory performance.

Here, we collected fMRI data while a group of younger and older adults learned images of faces and houses and subsequently performed an old/new recognition memory test. Using exploratory pattern similarity searchlight analyses, we looked for regions expressing high neural distinctiveness during memory retrieval as well as in encoding-retrieval reinstatement. Our key questions were whether we would find evidence of age-related neural dedifferentiation during retrieval and whether retrieval-related dedifferentiation contributes to senescent memory decline. We expected older adults to demonstrate reduced distinctiveness during retrieval, particularly in visual processing regions. We further predicted that lower distinctiveness during retrieval would be associated with poorer memory performance. Additionally, we compared our retrieval-related findings to encoding-retrieval reinstatement in order to understand whether age deficits in distinctiveness during retrieval extend beyond weakly reinstated encoding patterns.

## 2 Materials and Methods

Encoding and retrieval data from this project were previously reported in two papers (Kobelt et al., 2021; Pauley et al., 2022) that were later retracted by the authors due to a preprocessing error. For the retracted manuscripts as well as comparison reports with the corrected findings, please see https://osf.io/t8dpv/ and https://osf.io/7n3mz/.

### 2.1 Participants

Data were collected from a total of 76 healthy adults. Participants were recruited within two age groups: younger adults (18–27 years, *N* = 39) and older adults (64–76 years, *N* = 37). Two participants were excluded due to too much motion in the scanner (1 younger adult and 1 older adult), 3 were excluded due to memory performance below chance level (2 younger adults and 1 older adult), and 1 younger adult was excluded due to poor MRI data quality. The final sample consisted of 35 younger adults (*M* (*SD*) age = 22.3 (2.7) years, 16 females, 19 males) and 35 older adults (*M* (*SD*) age = 70.6 (2.4) years, 19 females, 16 males). Participants were screened via telephone for mental and physical illness, metal implants and current medications. Additionally, all older adults were screened using the Mini-Mental State Examination (Folstein et al., 1975) and all exceeded the threshold of 26 points. The study was approved by the ethics committee of the German Society for Psychological Research (DGPs) and written informed consent was obtained from each participant at the time of the study.

### 2.2 Stimuli

Stimuli were comprised of 300 grayscale images belonging to 3 different categories: 120 neutral faces (adapted from the FACES database; Ebner et al., 2010), 120 houses (some adapted from DC Park et al., 2004, and some obtained online), and 60 phase-scrambled images (30 faces and 30 houses, constructed from randomly selected face and house images) serving as control stimuli. An additional image from each category was selected to serve as target stimuli for the encoding target-detection task. All nontarget face and house images were randomly divided into 2 sets of 120 images (60 faces and 60 houses). One stimulus set was presented during both encoding and recognition (old images) and the other set was presented only during recognition (new images). The same stimulus sets were used for all participants.

### 2.3 Paradigm

The following paradigm was part of a larger study spanning two days of data collection. The present study focuses only on the face-house task, which comprised an incidental encoding phase and a surprise recognition test, both conducted inside the fMRI scanner on the same day with a delay of approximately 30 minutes (see Figure 1). The encoding phase consisted of 2 identical runs each with 9 stimulus blocks. Stimuli were randomly distributed into the blocks such that each block contained 20 images of a single category (faces, houses, or phase-scrambled) as well as the category’s corresponding target image. The block order was alternating and counterbalanced across participants, always starting with either a face or house block. The stimulus order within each block was pseudo-randomized with the condition that the target image was not presented in either the first 4 or last 4 trials of a block. Due to a technical problem, the same stimulus order was used for all participants who started with a face block and for 36 of the participants starting with a house block. In order to ensure the participants were paying attention to the stimuli, they were asked to perform a target-detection task in which they pressed a button when one of the 3 target images was presented. Prior to the encoding phase, participants completed 5 practice trials of each stimulus category, including each of the target stimuli, to verify that they understood the target-detection task. The nontarget training stimuli were excluded from the main experiment. Since the 2 encoding runs were identical, participants were exposed to each stimulus twice during the encoding phase. Stimuli were presented for 1200 ms and separated by a fixation cross with a jittered duration between 500 and 8000 ms. In total, the encoding phase lasted approximately 22 minutes.

**Figure 1.**
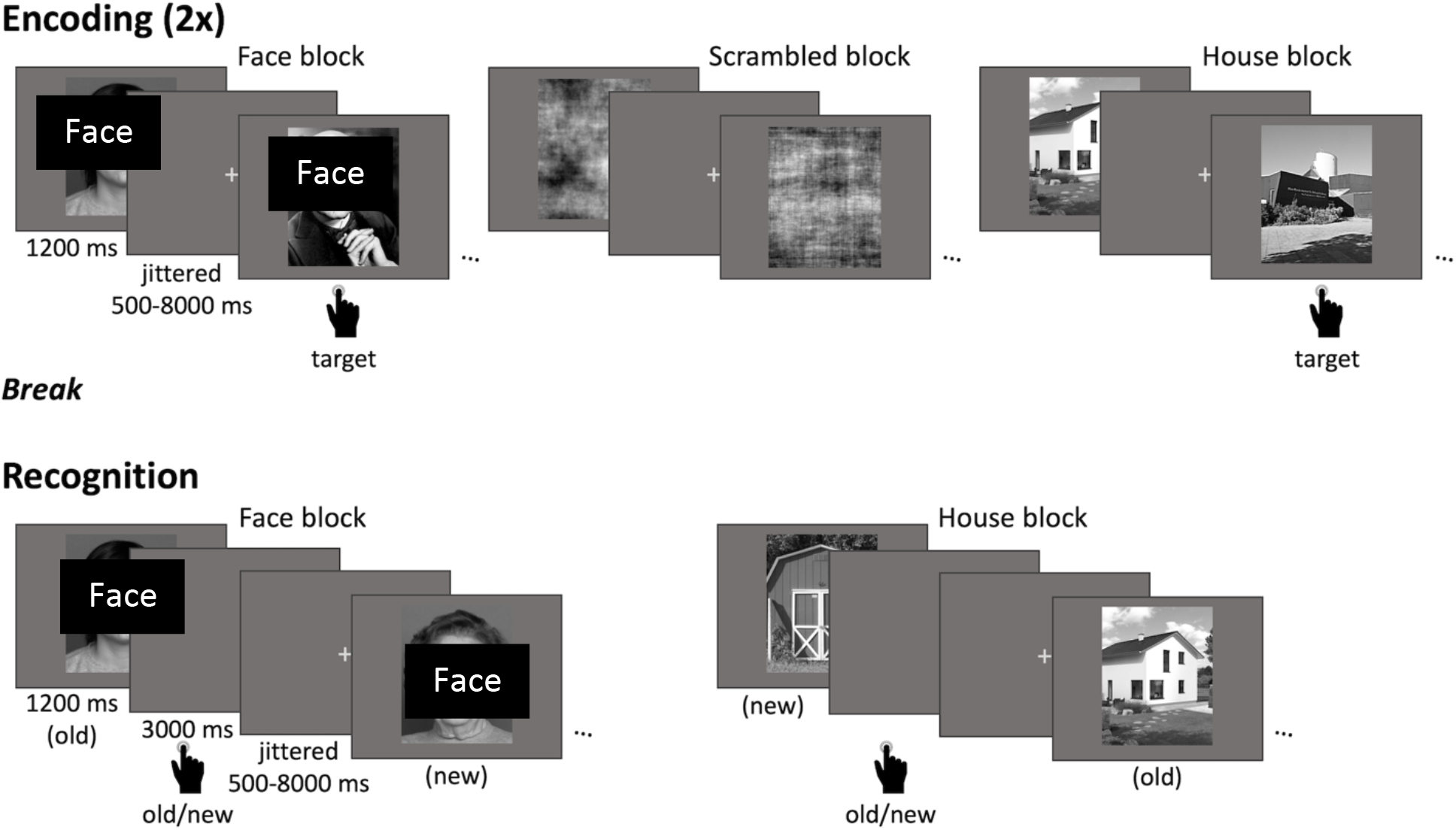
Face-house task design. This fMRI paradigm comprised an incidental encoding phase (top) and a surprise recognition test (bottom). During encoding, two identical runs of face, house, and phase-scrambled images were presented in a block design with 9 stimulus blocks each (3 alternating blocks from each stimulus category). Each block had 21 trials (20 exemplars of the respective category and 1 pre-learned target stimulus). Participants were instructed to press a button when a target stimulus appeared. During the recognition test, six alternating face and house blocks were presented with 40 trials each (20 old trials from encoding and 20 new trials). Participants indicated via button press whether each image was old or new. (Faces were made unidentifiable for the upload on Biorxiv.)

Following encoding, participants remained in the scanner briefly while structural scans were collected (see below). Then, they had a short break outside the scanner while they received instructions for the recognition test. They then returned to the scanner to complete the recognition test. The recognition test consisted of 6 blocks in total, alternating between 3 face and 3 house blocks and was divided into 2 runs of 3 blocks each. Each block contained 20 old images (seen during encoding) and 20 new images of the same stimulus category. For each trial, participants were asked whether the image was old or new, which they indicated via button press. The stimulus order was pseudo-randomized such that no more than 3 old or new images were presented consecutively. Due to a technical problem, the same stimulus order was used for 13 participants who started with a face block and 14 participants who started with a house block. Stimuli were presented for 1200 ms and followed by a gray screen for 3000 ms in which participants could give their response. Fixation crosses separated the trials with jittered durations between 500 and 8000 ms. In total, the recognition task lasted approximately 26 minutes.

### 2.4 fMRI data acquisition and preprocessing

Brain imaging was acquired with a Siemens Magnetom TrioTim 3T MRI scanner with a 32-channel head-coil. Functional images were collected using an echo planar imaging (EPI) sequence during both the encoding and recognition phases in 2 runs each. Each encoding run consisted of 270 volumes and each recognition run consisted of 372 volumes (voxel size = 3 x 3 x 3.3 mm^3^; TR = 2 s; TE = 30 ms). The first three volumes of each run were dummy volumes and were excluded prior to preprocessing. Following the encoding phase, a T1-weighted (T1w) magnetization prepared rapid acquisition gradient echo (MPRAGE) pulse sequence image was acquired (voxel size = 1 x 1 x 1 mm^3^; TR = 2.5 ms; TE = 4.77 ms; flip angle = 7°; TI = 1.1 ms). Additionally, turbo spin-echo proton density images (PDs), diffusion tensor images (DTIs), and fluid attenuation inversion recovery images (FLAIRs) were collected, but not included in the following analyses. Experimental stimuli, which participants viewed via a mirror mounted on the head-coil, were projected using the Psychtoolbox (Psychophysics Toolbox) for MATLAB (Mathworks Inc., Natick, MA).

MRI data were organized according to the Brain Imaging Data Structure (BIDS) specification (Gorgolewski et al., 2016) and preprocessed using *fMRIPrep* (version 1.4.0; Esteban et al., 2019) with the default settings. The T1w image was corrected for intensity nonuniformity, skull-stripped, and spatially-normalized to the *ICBM 152 Nonlinear Asymmetrical template version 2009c* through nonlinear registration. Functional images were motion-corrected, slice-time corrected, and co-registered to the normalized T1w reference image. Finally, functional images were resampled to 2 mm isotropic voxels in order to enhance the signal-to-noise ratio (Dimsdale-Zucker & Ranganath, 2018).

### 2.5 Behavioral data analyses

Behavioral analyses were performed using custom MATLAB scripts. Recognition memory performance (*Pr*) was measured as the difference between the hit rate (proportion of correctly identified old stimuli) and the false alarm rate (proportion of new stimuli incorrectly identified as old stimuli; Snodgrass & Corwin, 1988). An independent-samples *t*-test was used to assess age differences in memory performance and dependent-samples *t*-tests were used to determine whether memory performance differed between face and house stimuli and whether memory performance exceeded chance level. Age differences in response bias were assessed with independent-samples *t*-tests comparing the hit rates and false alarm rates across age groups.

### 2.6 Pattern similarity searchlight analyses

In order to perform pattern similarity analyses, a generalized linear model (GLM) was performed for each trial in both encoding and recognition, including one trial-specific regressor, one regressor for all other trials within the same run, and six motion regressors (Mumford et al., 2012). Trial regressors were modeled as 1.2 s duration boxcar functions convolved with a canonical hemodynamic response function. Pattern similarity analyses were based on the resulting β weights for each trial. Pattern similarity was only assessed between trials from different runs to control for time-dependent correlations in the hemodynamic responses (Dimsdale-Zucker & Ranganath, 2018) and was measured as Fisher *z*-transformed Pearson correlations. Searchlight similarity analyses were conducted using modified scripts from the MATLAB toolbox for representational similarity analysis (Nili et al., 2014) and with 8-mm-radius spherical searchlights.

Several searchlight similarity measures were computed (see Figure 2). First, in order to identify brain regions demonstrating high category specificity across all participants *during memory recognition*, we compared within-category similarity to between-category similarity for both faces and houses separately. For each stimulus, within-category similarity was calculated as the averaged across-voxel correlation of the activity pattern in response to the stimulus to the activity patterns in response to all other stimuli from the same category (e.g., mean similarity of the activity in response to a face stimulus to that of all other face stimuli). For each participant, within-category similarity was then averaged across all stimuli within each category. Between-category similarity was calculated as the averaged across-voxel correlation of the stimulus’ activity pattern to the activity patterns of all stimuli from the other category (e.g., mean similarity of the activity in response to a face stimulus to that of all house stimuli). Between-category similarity was then averaged across all stimuli for each participant. Within- and between-category similarity were assessed in a searchlight centered on each voxel in the brain and the difference was calculated, resulting in a whole-brain map of category specificity for faces and a whole-brain map of category specificity for houses in each participant.

**Figure 2.**
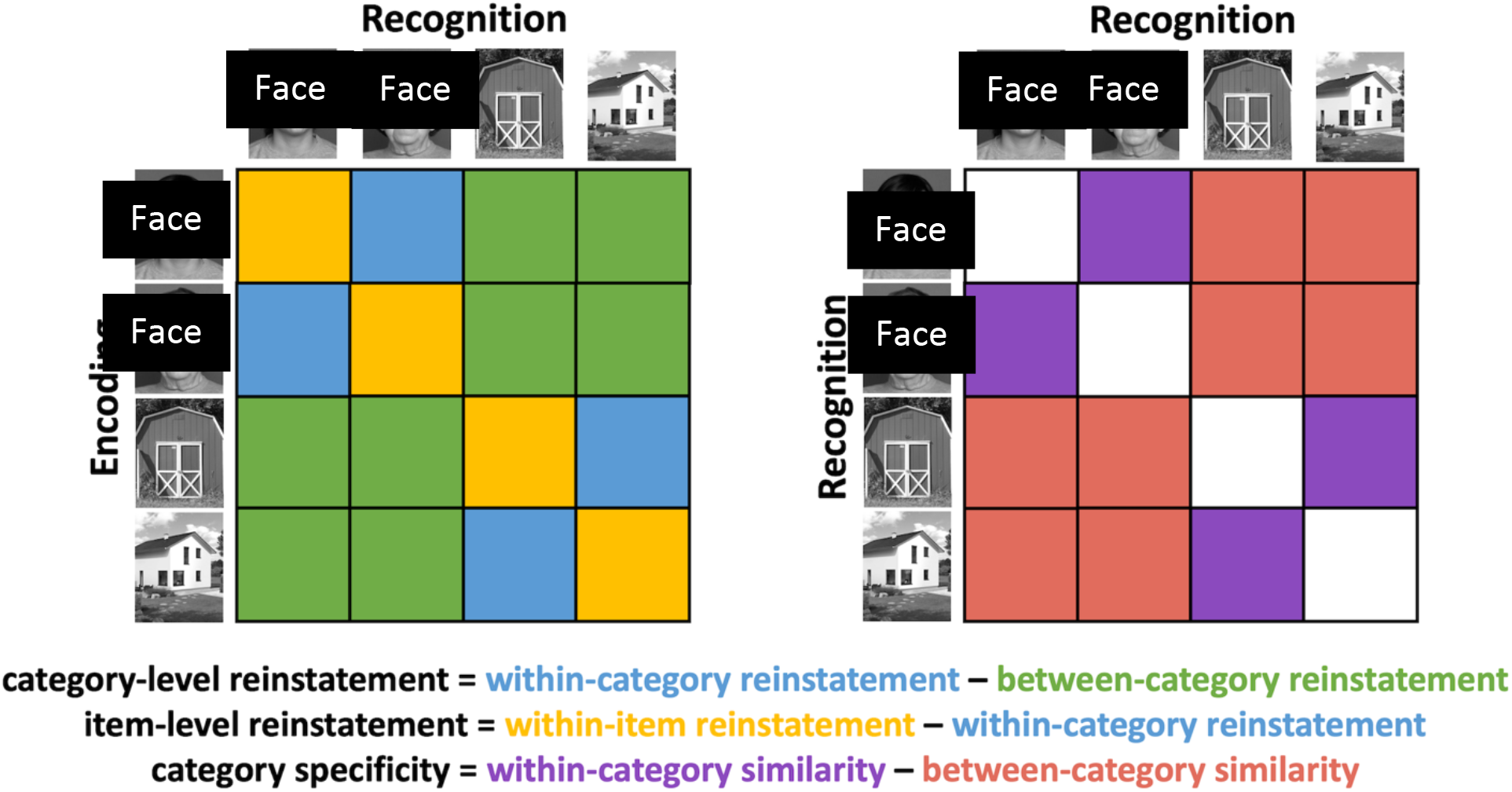
Illustration of searchlight similarity measures calculated during encoding-retrieval reinstatement (left) and during recognition (right).

Next, we were interested in searching for brain regions exhibiting *encoding-retrieval reinstatement* both at the category level and at the individual stimulus level. In order to assess category-level reinstatement, we compared within-category reinstatement to between-category reinstatement. Within-category reinstatement was calculated as the mean pattern similarity of all stimuli from a given category during encoding to all stimuli from the same category during recognition. Within-item reinstatement correlations (i.e., the similarity of a given stimulus’ activity pattern during encoding to the activity pattern of the same stimulus during recognition) were excluded from the measure of within-category reinstatement. Between-category reinstatement was similarly calculated as the mean pattern similarity of all stimuli from a given category during encoding to all stimuli from the other category during recognition. Within- and between-category reinstatement were then averaged across all stimuli in each searchlight and the difference in each voxel was computed, resulting in a whole-brain map of category-level reinstatement for each participant. A whole-brain map of item-level reinstatement was created for each participant by calculating the voxel-wise difference between within-item reinstatement and within-category reinstatement. For both category- and item-level reinstatement, similarity values were calculated between recognition and each encoding run individually, then averaged across encoding run.

For each searchlight similarity measure, nonparametric, cluster-based, random permutation analyses adapted from the FieldTrip toolbox (Oostenveld et al., 2011) were used to identify brain regions demonstrating significant effects across all participants (e.g., for the high category specificity measures, regions demonstrating greater within-category similarity than between-category similarity). First, dependent-samples *t*-tests were conducted within each voxel. Adjacent voxels with significance values lower than a threshold of *p* < 0.005 were grouped into clusters. The sum of all *t* statistics of the voxels included in each cluster was defined as the cluster test statistic. The Monte Carlo method was used to determine whether a cluster was significant by comparing the cluster test statistic to a reference distribution of *t* statistics across 1000 permutations. Each *t* statistic in the reference distribution was created by randomly reallocating the two conditions and calculating the cluster test statistic based on this random reallocation. Clusters were considered significant under a threshold of *p* < 0.05 and if they contained at least 10 voxels.

### 2.7 Assessing age differences in searchlight similarity analyses

We were also interested in searching for brain regions demonstrating age differences in each of our searchlight similarity measures. For our age comparison analyses, we limited the search space to the regions identified during the whole-group analyses (i.e., age differences were only assessed in regions demonstrating an effect across all participants). Since some of the clusters identified during the whole-group analyses were fairly large (>8000 voxels), we again used nonparametric, cluster-based, random permutation analyses to search these clusters for regions exhibiting age differences. First, independent-samples *t* tests were conducted within each voxel comparing younger and older adults on each searchlight similarity measure described previously. Adjacent voxels with significance values lower than a threshold of *p* < 0.005 were grouped into clusters and the sum of all *t* statistics of the voxels included in each cluster was defined as the cluster test statistic. The Monte Carlo method was again used to determine whether a cluster was significant. In this case, the reference distribution was created by removing the younger and older adult labels and randomly assigning participants to each age group across 1000 permutations.

### 2.8 Relating searchlight similarity measures to memory performance

We further assessed the relationship between our searchlight similarity measures and interindividual differences in memory performance. Therefore, permutation testing was performed again for each cluster identified during the whole-group analyses. For this, regressions were conducted predicting memory performance from the searchlight similarity measure in each voxel (i.e., recognition category specificity, category-level reinstatement, item-level reinstatement). As described previously, adjacent voxels below the threshold of *p* < 0.005 were grouped into clusters, the cluster test statistic was calculated, and the Monte Carlo method determined the significance of each cluster across 1000 permutations. In order to derive the correlation coefficient for significant clusters to better understand the relationships, we averaged the respective searchlight similarity measure across all voxels within the cluster for each participant and correlated this with *Pr* across participants using Pearson partial correlations controlled for group differences (these can be found in Table 3). Clusters identified during face analyses were correlated with *Pr* for faces and clusters identified during house analyses were correlated with *Pr* for houses.

Additionally, we were interested in whether recognition category specificity or category-level reinstatement was better at tracking interindividual variability in memory performance. To this end, we computed two linear model comparisons predicting *Pr*. For the first model comparison, one linear model was computed using age and recognition category specificity (i.e., Pr ∼ Age* RecSpec) and the other model added the interaction between age and category-level reinstatement (i.e., Pr ∼ Age* RecSpec + Age*ReinSpec). The second model comparison included age and category-level reinstatement as predictors in one linear model and additionally the interaction between age and recognition category specificity in the other model. In this way, we were able to determine whether recognition category specificity or category-level reinstatement better explained memory-related variance. The models were compared using the *anova()* function in *R*.

### 2.9 Trial-wise mixed effects analyses

Recent work has pointed to the significance of both reinstatement as well as hippocampal activity in predicting within-person variability in memory performance in both younger and older adults (Trelle et al., 2020; Hill et al., 2021). In order to investigate this further, we performed generalized linear mixed-effects models to determine whether item-level reinstatement specificity or trial-wise hippocampal activity were successful predictors of intraindividual differences in memory performance. To measure trial-wise hippocampal activity, we used a bilateral hippocampal mask defined by the automatic anatomic labeling atlas. For each participant, we averaged across all beta weights within the hippocampal mask for every trial and z-transformed across trials within each participant. Since we observed age differences in response bias, we also included a participant-wise measure of response bias in the model defined as FAR/(1 – (HR – FAR)), where FAR is the false alarm rate and HR is the hit rate (Snodgrass & Corwin, 1988). We performed a separate model for each cluster resulting from our item-level reinstatement searchlight similarity analyses. For each model, binary recognition memory outcomes (hit or miss) were used as the dependent variable and age, response bias, hippocampal activity, and item-level reinstatement (operationalized as the difference between the similarity of an item to itself across encoding and recognition and the mean similarity of the same item to all other items within the same stimulus category) were used as independent variables. The interactions between age and hippocampal activity as well as age and item-level reinstatement were included in the model. The model was analyzed using the *R* function *glmer* from the lme4 package with the following formula: Memory ∼ Age + RespBias + Hipp + ItemRein + Age*Hipp + Age*ItemRein + (1 + Hipp + ItemRein | Subject).

## 3 Results

### 3.1 Behavioral results

We first checked for age differences in memory performance (i.e., *Pr* = hit rate – false alarm rate). Memory performance did not differ between age groups (*M*_younger_ = 0.24, *SD*_younger_ = 0.12, *M*_older_ = 0.19, *SD*_older_ = 0.12, *t*(68) = 1.79, *p* = 0.08) and memory performance exceeded chance level in both younger (*t*(34) = 11.88, *p* < 0.001) and older adults (*t*(34) = 9.25, *p* < 0.001). Additionally, memory performance did not differ between face and house stimuli in either younger (*t*(34) = -0.88, *p* = 0.38) or older adults (*t*(34) = -1.60, *p* = 0.12). Older adults demonstrated a strong response bias, responding “old” more often than younger adults to both old stimuli (*M*_younger_ = 0.50, *SD*_younger_ = 0.14, *M*_older_ = 0.60, *SD*_older_ = 0.13, *t*(68) = -3.27, *p* = 0.002) and new stimuli (*M*_younger_ = 0.26, *SD*_younger_ = 0.11, *M*_older_ = 0.42, *SD*_older_ = 0.13, *t*(68) = - 5.42, *p* < 0.001).

### 3.2 Category specificity during recognition and reinstatement in occipital and ventral visual regions

Many studies have presented evidence for selective cortical reinstatement in sensory cortices associated with the encoded stimuli (for review, see Danker & Anderson, 2010). However, few studies have considered category specificity explicitly during memory retrieval, potentially missing the full picture of neural distinctiveness during retrieval. Therefore, we first searched for regions exhibiting greater within-category similarity than between-category similarity in the whole sample of participants for both faces and houses during recognition. Our searchlight similarity analysis revealed four clusters demonstrating category specificity for faces in ventral visual, temporal, and frontal regions during recognition (*p*s < 0.05; see Table 1 and Figure 3). We additionally identified one large cluster demonstrating strong category specificity for houses in ventral visual and occipital regions (*p* < 0.001).

**Figure 3.**
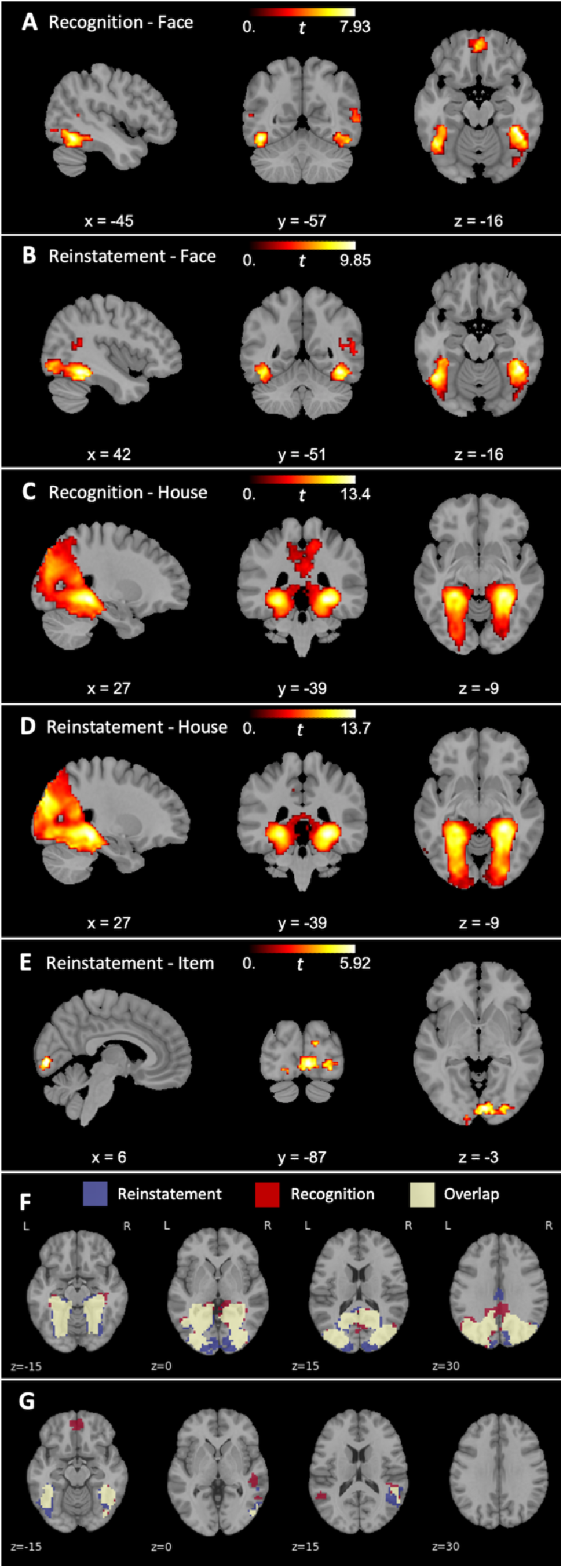
Regions demonstrating category specificity during recognition and reinstatement for both faces (A, B) and houses (C, D). Regions demonstrating item-level reinstatement specificity for houses (E). Overlapping voxels in category specificity for houses (F) and faces (G) between recognition and reinstatement. Blue = voxels only identified during category-level reinstatement; red = voxels only identified during recognition; yellow = voxels overlapping between recognition and reinstatement.

**Table 1.**
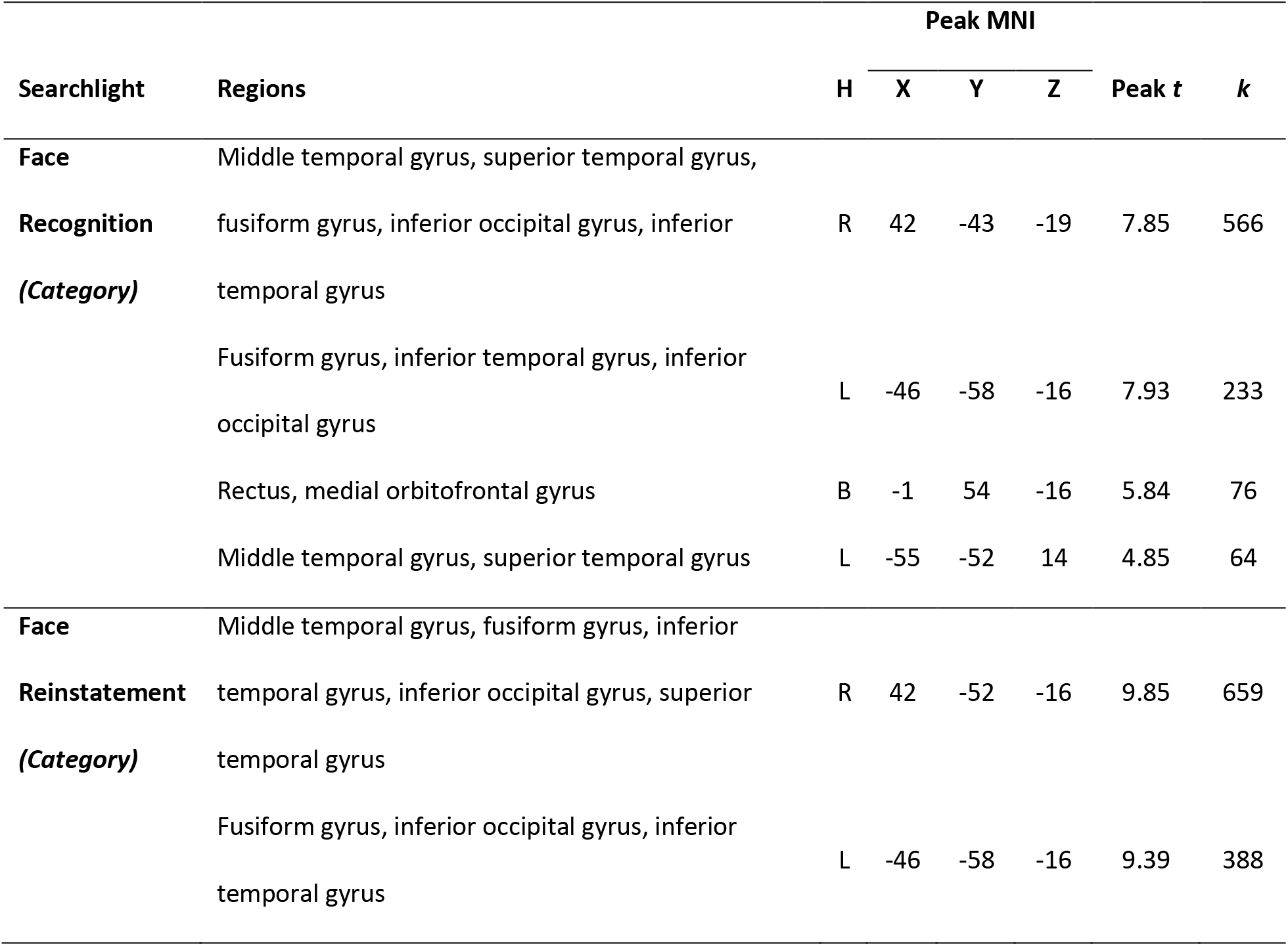
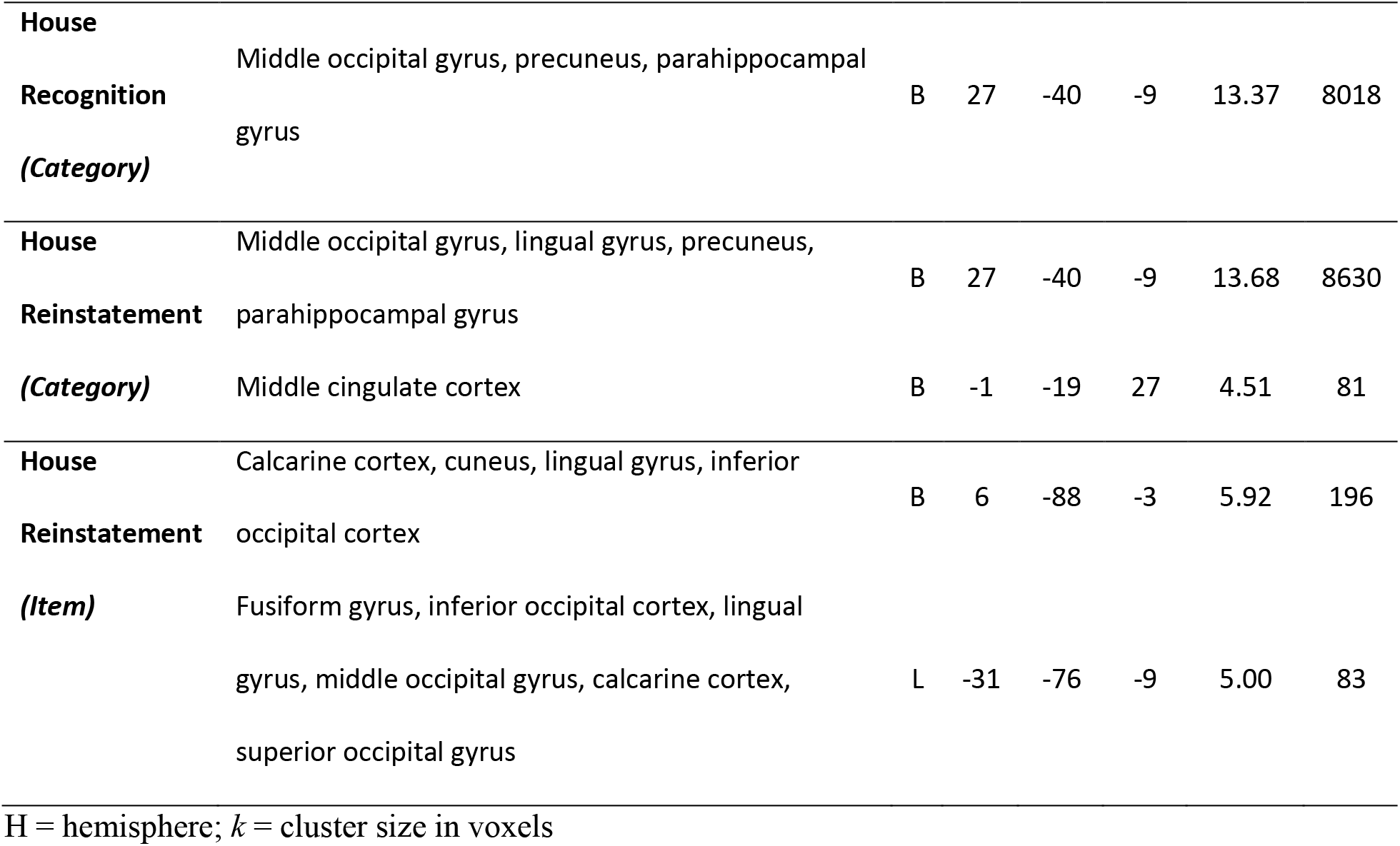
Clusters identified by searchlight similarity analyses revealing high category specificity during recognition and reinstatement

Next, we searched for regions demonstrating encoding-retrieval reinstatement at the category level for faces and houses. We found two clusters demonstrating strong category-level reinstatement for faces in temporal and occipital regions (*p*s < 0.001; see Table 1 and Figure 3) and two clusters demonstrating strong category-level reinstatement for houses in occipital regions (*p* < 0.04). Importantly, there was a high degree of overlap between category-specific regions identified during recognition and reinstatement with 6827 voxels identified during recognition and reinstatement overlapping for houses and 640 voxels overlapping for faces (see Figure 3). Thus, specificity during recognition is largely dependent on reinstated encoding specificity, though with some variation.

We also searched for regions demonstrating encoding-retrieval similarity at the item level for both faces and houses. These item-level reinstatement searchlight similarity analyses yielded two clusters in occipital regions for houses (*p*s < 0.02; see Table 1 and Figure 3), but no significant clusters for faces (*p*s > 0.08).

### 3.3 Age differences in category specificity during recognition and category-level reinstatement

Previous findings reveal clear age deficits in reinstatement specificity (St-Laurent et al., 2014; Johnson et al., 2015; Abdulrahman et al., 2017; Bowman et al., 2019; Trelle et al., 2020; Hill et al., 2021), particularly in occipital and temporal regions, but age deficits in retrieval-related specificity are relatively less documented (St-Laurent et al., 2014; Johnson et al., 2015). Accordingly, we tested for age differences in category specificity, limiting the search space to regions identified by the whole-group analyses. Cluster permutation testing revealed two clusters demonstrating age differences in category specificity for houses during recognition (*p*s < 0.001) and three clusters demonstrating age differences in category-level reinstatement for houses (*p*s < 0.05) in bilateral ventral visual cortices (see Table 2 and Figure 4). Younger adults exhibited greater category specificity than older adults in all clusters. No age differences were identified in category specificity for faces during either recognition or reinstatement (all clusters were smaller than 10 voxels). Furthermore, no age differences were identified in item-level house specificity. Although there was substantial overlap in age differences between retrieval and reinstatement, age differences during retrieval were spatially more widespread than age differences during reinstatement (see Figure 4). Age differences in the size of clusters during retrieval spanned 61,271 mm^3^. Of this volume, 41,432 mm^3^ (or 68%) were shared with age differences in reinstatement, while 19,839 mm^3^ were unique to retrieval. By contrast, age differences in reinstatement spanned 47,936 mm^3^, with 41,432 mm^3^ (or 86%) overlapping with age differences in recognition and 6,504 mm^3^ reflecting age differences unique to reinstatement. Hence, age differences in category specificity are spatially more extended during retrieval processing, while age differences during reinstatement are more localized.

**Figure 4.**
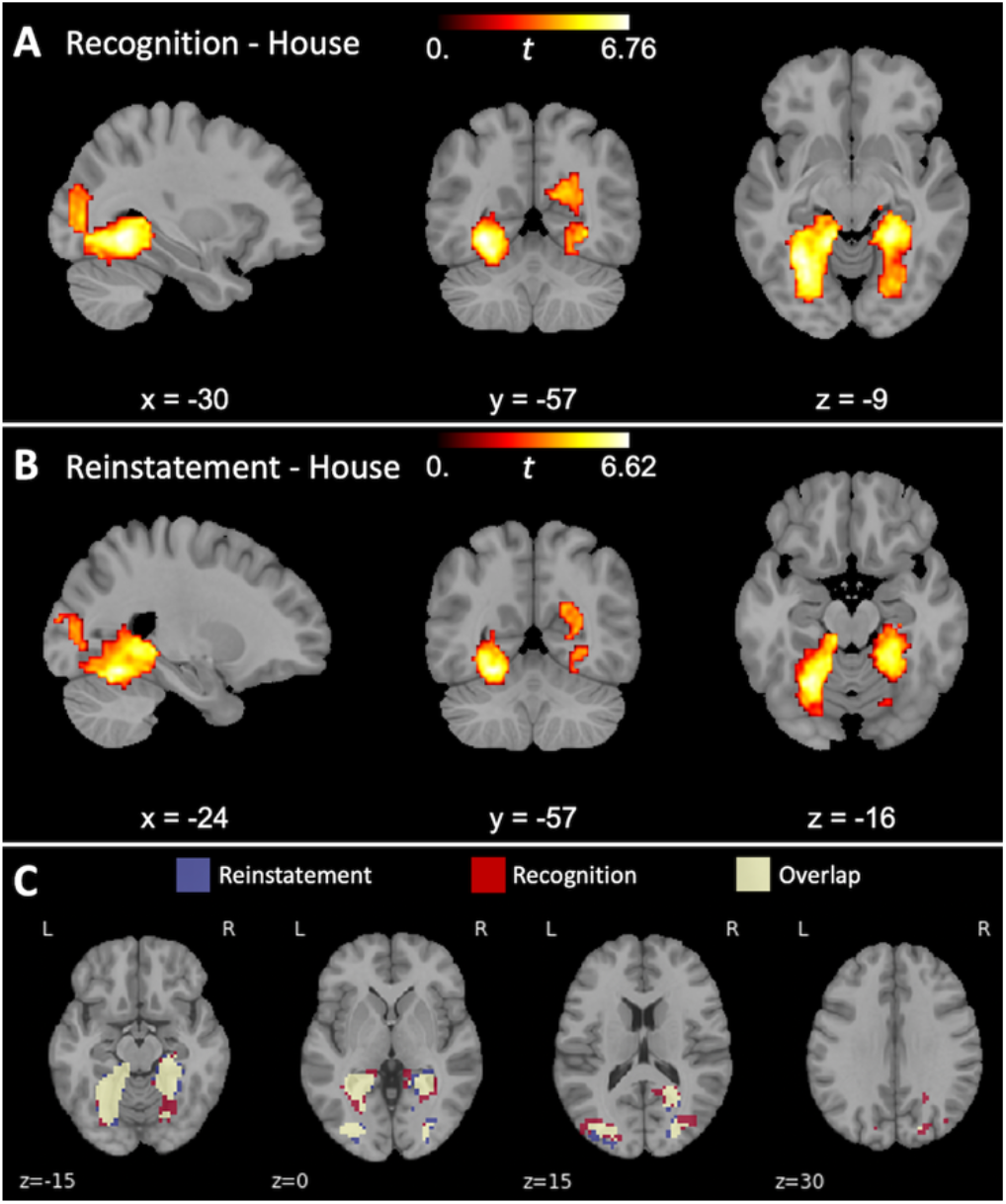
Age differences in category specificity for houses. Younger adults demonstrated greater category specificity (within-category similarity > between-category similarity) than older adults during both recognition (A) and reinstatement (B) for house stimuli. Overlapping voxels in age differences in category specificity for houses between recognition and reinstatement (C). Blue = voxels only identified during category-level reinstatement; red = voxels only identified during recognition; yellow = voxels overlapping between recognition and reinstatement.

**Table 2.**
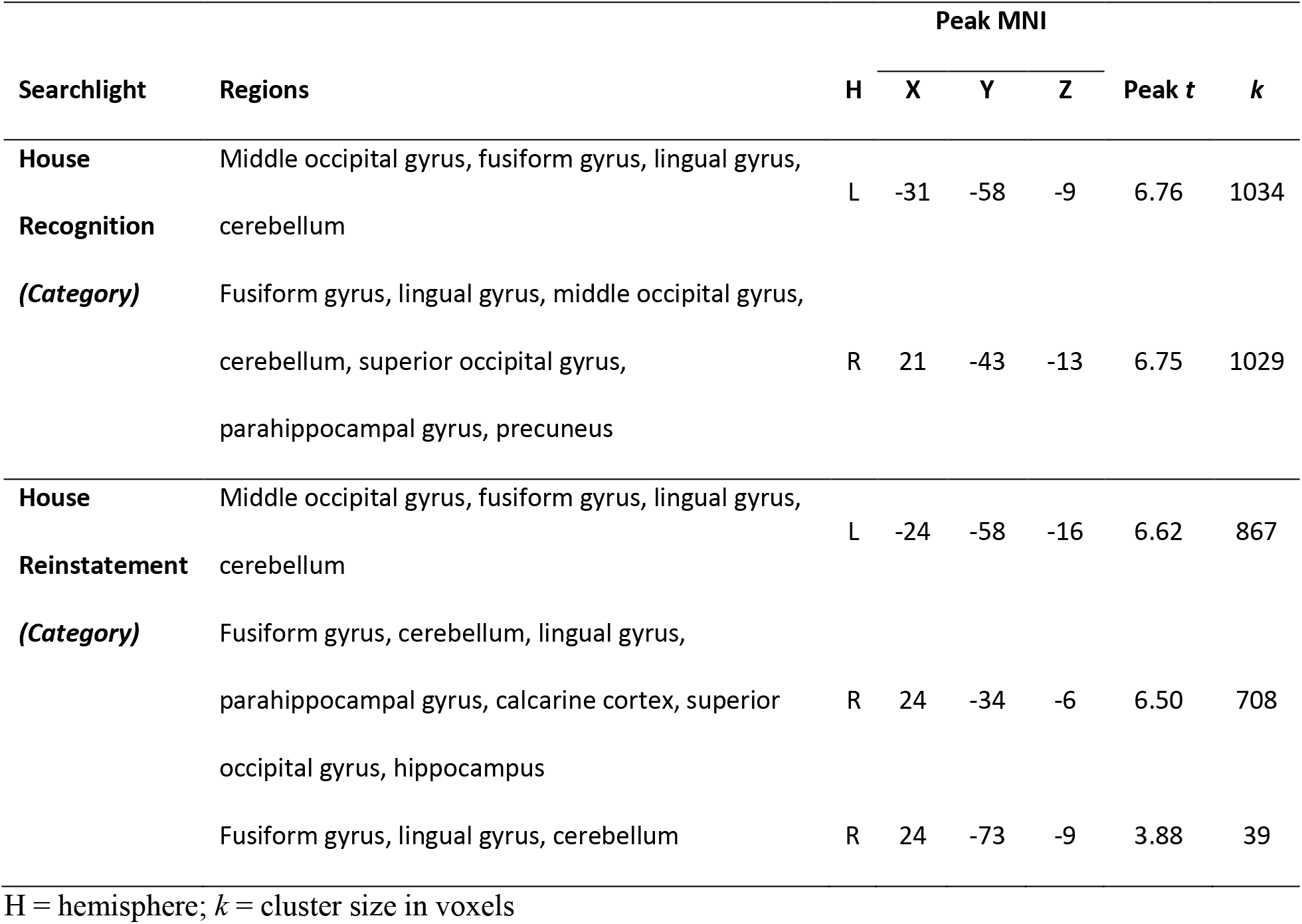
Clusters revealing greater category specificity in younger adults than in older adults

### 3.4 Category-level reinstatement predicts interindividual differences in memory performance to a greater spatial extent than recognition

Here, we asked whether category specificity in the regions identified by the whole-group analyses was linked to memory performance. In temporal cortices, we identified two clusters demonstrating a positive relationship between memory performance and category-level reinstatement for faces (*p*s < 0.001; see Table 3 and Figure 5). In occipital and ventral visual regions, we also found a cluster revealing a positive relationship between memory performance and category-level reinstatement for houses (*p* < 0.001). Two clusters were also identified revealing a positive relationship between memory performance and category specificity during recognition for houses (*p*s < 0.001). However, we did not identify any regions demonstrating a relationship between memory performance and face specificity during recognition (all clusters had fewer than 10 voxels). Furthermore, we did not identify any clusters revealing a relationship between memory performance and item-level reinstatement specificity for houses. Therefore, only category-level distinctiveness was positively associated with interindividual differences in memory performance, particularly for house stimuli. Crucially, category reinstatement specificity tracked memory to a greater spatial extent than recognition specificity (see Figure 5). The size of the clusters revealing an association between memory performance and category specificity during retrieval spanned 10,514 mm^3^. Of this volume, 9,950 mm^3^ (or 95%) were shared with the correlation between memory and reinstatement specificity, while 564 mm^3^ were unique to retrieval. By contrast, the size of the clusters showing an association between memory performance and category-level reinstatement spanned 71,993 mm^3^, with 9,950 mm^3^ (or 14%) overlapping with the correlation between memory and recognition specificity and 62,043 mm^3^ reflecting a relationship unique to reinstatement. Therefore, we found that both category specificity during recognition and category-level reinstatement were associated with memory performance, such that participants with more specific neural representations also tended to have better memory performance. However, the volume of the regions identified as correlating with memory performance was greater for category-level reinstatement than recognition category specificity.

**Figure 5.**
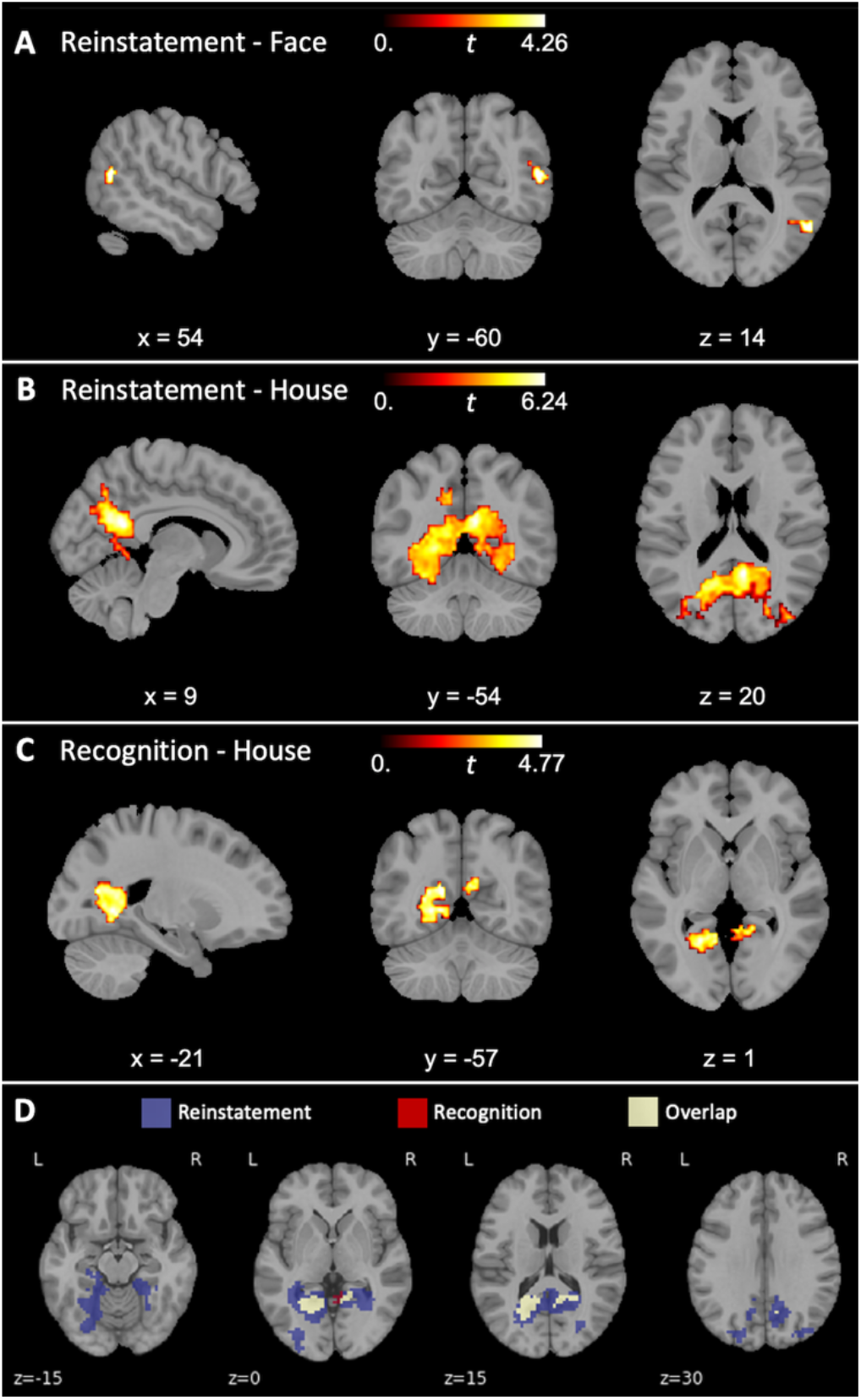
Regions indicating a relationship between category-level specificity and memory performance for reinstated faces (A), reinstated houses (B) and houses during recognition (C). Overlapping voxels in the correlation between category specificity and memory performance for houses between recognition and reinstatement (D). Blue = voxels only identified during category-level reinstatement; red = voxels only identified during recognition; yellow = voxels overlapping between recognition and reinstatement.

**Table 3.**
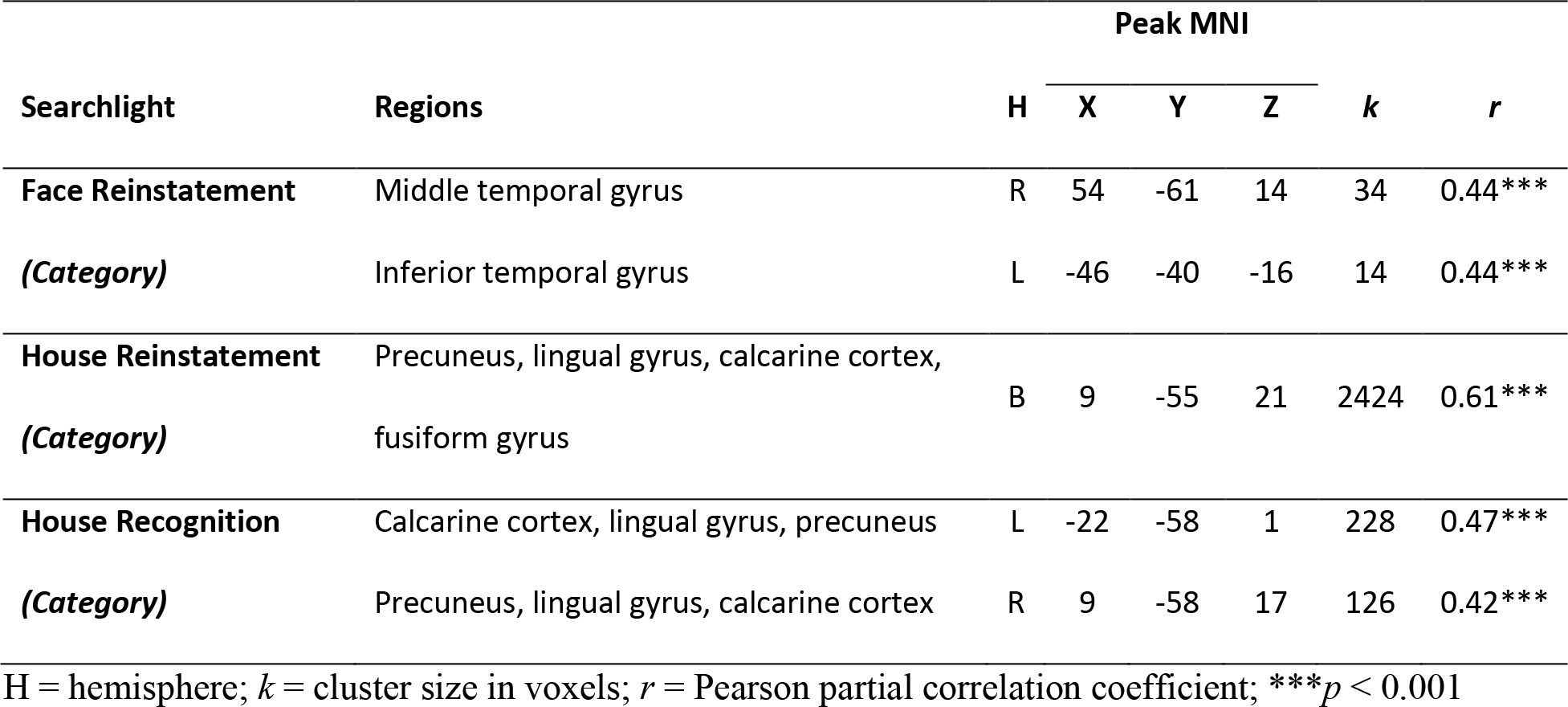
Clusters in which category specificity correlates with memory performance

### 3.5 Category-level reinstatement tracks interindividual variability in memory performance better than recognition category specificity

We additionally investigated whether either recognition category specificity or category-level reinstatement was better at explaining interindividual differences in memory performance using two linear model comparisons. The first model comparison revealed that adding reinstatement as a predictor improved the model fit on memory performance (*R^2^* = 0.41) as compared with recognition specificity (*R^2^* = 0.24; *F*(64) = 9.64, *p* < 0.001). However, the second model comparison revealed that adding recognition specificity did not improve the model fit on *Pr* (*R^2^* = 0.41) as compared with reinstatement (*R^2^* = 0.39; *F*(64) = 1.21, *p* = 0.31). These findings suggest that between-participant variability in memory performance is best explained by category-level reinstatement as compared with recognition category specificity.

### 3.6 Intraindividual variability in memory performance covaries with item-level reinstatement

In the following, we were interested in whether the ability to reinstate item-level encoding information or hippocampal activity during retrieval were associated with within-person memory outcomes. Our item-level reinstatement searchlight similarity analysis for houses revealed two significant clusters: one primarily located in the calcarine cortex and the other located in and around the fusiform cortex. For each cluster, we performed a general linear mixed effects model in order to test whether binary memory response outcome (hit or miss) could be predicted by trial-wise item reinstatement or trial-wise retrieval-related hippocampal activity. Our results revealed a significant effect of response bias in both clusters (calcarine cluster: log odds = 27.27, 95% CI [16.72–44.46]; fusiform cluster: log odds = 26.51, 95% CI [16.09–43.69]), showing that a higher bias for “old” responses was related to a higher hit rate. We found that trial-wise item reinstatement predicted memory outcome in both clusters (calcarine cluster: log odds = 2.37, 95% CI [1.38–4.05]; fusiform cluster: log odds = 1.68, 95% CI [0.99–2.84]), demonstrating that greater item-level reinstatement was beneficial for memory performance. We additionally found a trend interaction between age and item reinstatement in the calcarine cluster (log odds = 0.45, 95% CI [0.19– 1.03]), indicating that greater item reinstatement benefited memory performance in younger adults (see Figure 6; *t* = 3.39, *p* < 0.001, *M*_hit_ = 0.041, *SD*_hit_ = 0.178, *M*_miss_ = 0.015, *SD*_miss_ = 0.170), but not in older adults (*t* = 0.12, *p* = 0.90, *M*_hit_ = 0.031, *SD*_hit_ = 0.145, *M*_miss_ = 0.030, *SD*_miss_ = 0.145). No other fixed effects or interactions reached significance (see Table 4). Thus, item reinstatement, but not hippocampal activity, was predictive of intraindividual variability in memory performance. We additionally identified an age-related interaction, such that younger adults’ memory benefited from precise item reinstatement, but older adults’ memory did not.

**Figure 6.**
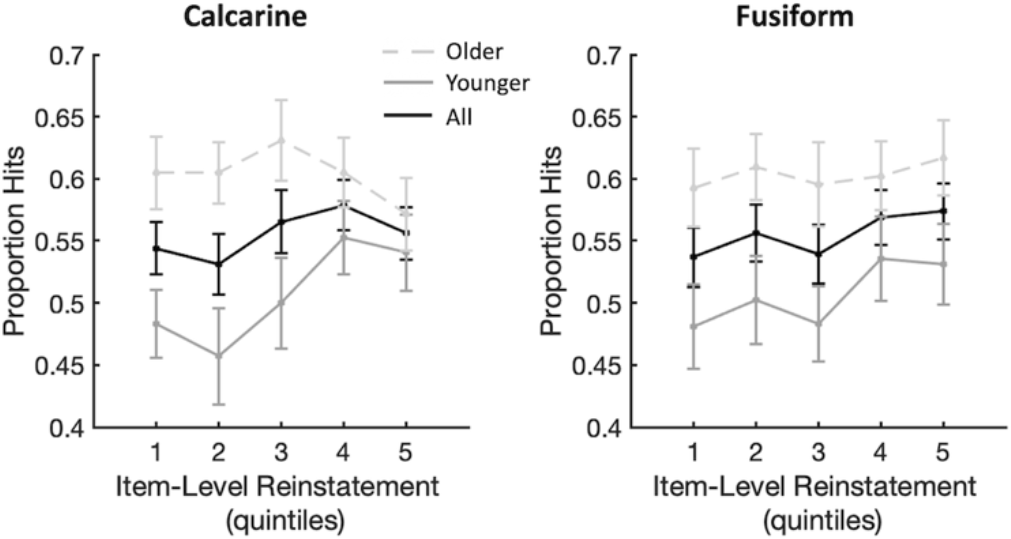
Increasing item-level reinstatement predicts a higher proportion of hits in the calcarine cluster (left) and fusiform cluster (right). Older adults are displayed with a dashed line, younger adults are displayed with a solid gray line, and an across-group average is displayed in black. For visualization, memory outcome was binned into quintiles according to trial-wise item reinstatement in each participant. Within each age group, the proportion of hits was averaged across participants. Error bars indicate ±1 standard error.

**Table 4.**
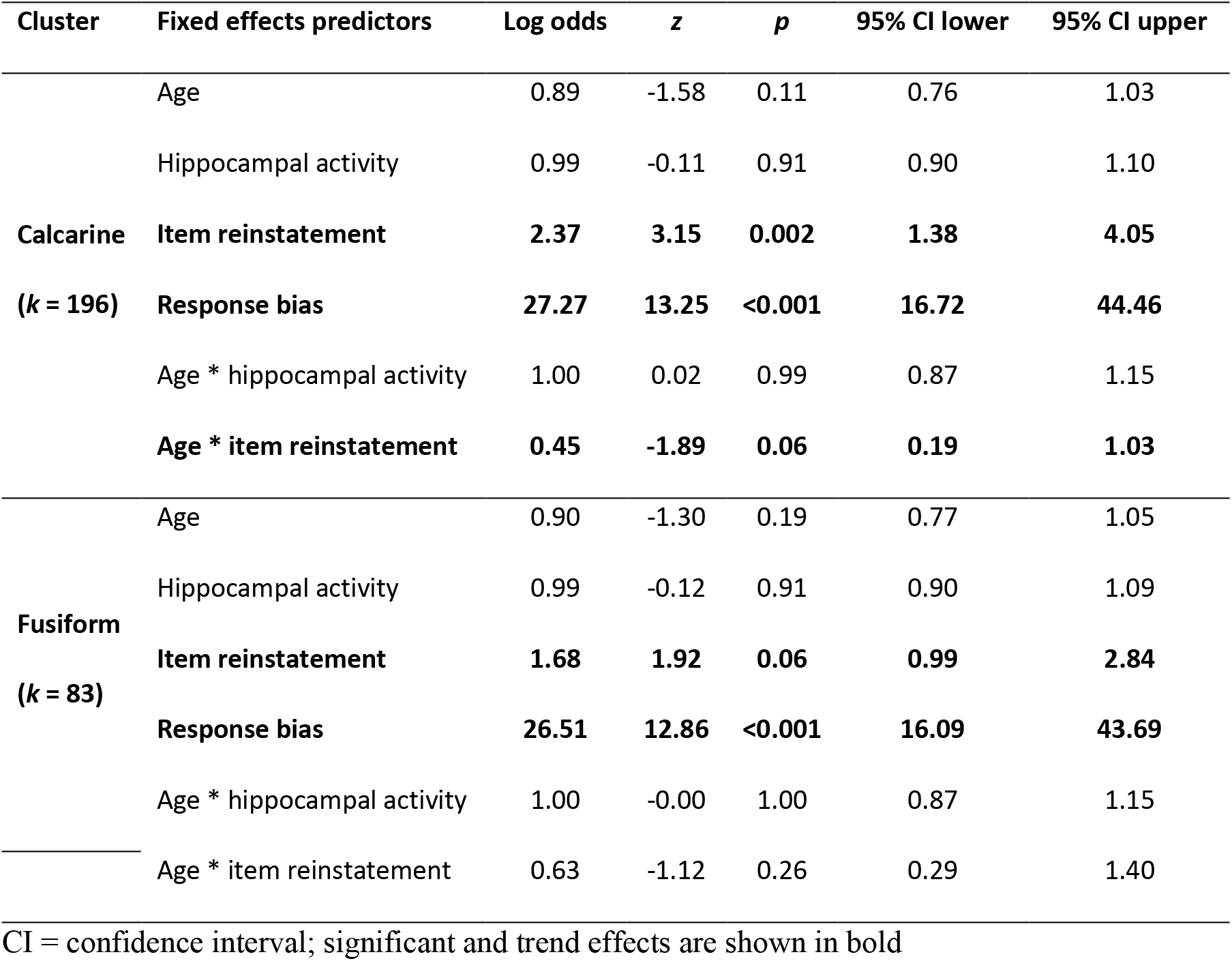
Results of general linear mixed effects models predicting trial-wise recognition memory accuracy

## 4 Discussion

The present study implemented exploratory multivariate pattern similarity searchlight analyses in order to investigate the influence of age-related neural dedifferentiation on category-sensitive neural representations during retrieval and encoding-retrieval reinstatement. Neural distinctiveness, reflected in less similar neural representations of items between different stimulus categories as compared to items within the same stimulus category, was observed during retrieval and reinstatement for both face and house stimuli. Importantly, younger adults demonstrated more distinctive (house) representations than older adults, in line with the age-related neural dedifferentiation hypothesis (Li et al., 2001; for review, see Koen & Rugg, 2019). The specificity of reinstated categorical representations was linked to better memory performance, providing evidence supporting the cortical reinstatement hypothesis (Danker & Anderson, 2010). Using a recognition paradigm, we were able to compare retrieval-related neural patterns of distinctiveness with reinstated encoding patterns and found that both overlapped highly. Despite this congruency, age differences in neural distinctiveness between younger and older adults were spatially more widespread during retrieval than during reinstatement. At the same time, reinstated neural patterns were more strongly linked to memory performance than retrieval-related distinctiveness. Thus, our findings suggest that while age-related neural dedifferentiation is more prominent during memory retrieval, the specificity with which neural patterns are reinstated seems to be more important for successful memory processing.

First and foremost, this study contributes to the scant literature on age-related neural dedifferentiation during memory retrieval. While many studies have highlighted the importance of representational distinctiveness during encoding (Zheng et al., 2018; Koen et al, 2019; Srokova et al., 2020; Hill et al., 2021), a recent review discussed how imprecise neural signaling during retrieval may lead to further memory impairment in older adults (Sander et al., 2021). In this review, Sander and colleagues (2021) highlighted how age differences both in mechanisms and neural structures supporting consolidation and retrieval amplify age differences already observed during encoding, resulting in poor memory quality. Therefore, our findings that reduced neural distinctiveness persists in older adults across memory stages reveal yet another hurdle for memory accuracy in older age.

Our focus on neural dedifferentiation exclusively during memory retrieval, as opposed to encoding-retrieval reinstatement, was motivated by recent reports of spatial and representational transformations in information processing between encoding and retrieval (Xiao et al., 2017; Favila et al., 2018; Srokova et al., 2022). These studies showed that, during retrieval, representations of encoding-related content can be found in regions other than those identified during encoding. For example, peaks of neural activation during retrieval have been found to be located more anteriorly compared to peaks of neural activation during encoding, termed the anterior shift (Bainbridge et al., 2021; for review, see Favila et al., 2020). The magnitude of this anterior shift has been reported to be larger in older adults than in younger adults (Srokova et al., 2022). These findings suggested that age differences in categorical representations might be salient outside reinstated encoding patterns. While we observed overlapping effects of dedifferentiation between recognition and reinstatement, age deficits in specificity during retrieval in our recognition paradigm clearly extended beyond poor reinstatement specificity. These age deficits during retrieval were limited primarily to occipital and ventral visual regions. Since we used a recognition task in the present study showing old and new stimuli during the retrieval phase, the observed age effects in these regions may be closely related to similar effects observed during encoding, and thus may mostly reflect an impairment in active perceptual (re-)processing during recognition. It is important to note that the type of retrieval task plays a role in which brain areas are recruited. Accordingly, a different retrieval task may also recruit category-specific representations in areas outside visual processing regions and reveal age differences therein. For example, in a recent study revealing encoding-retrieval transformation, Favila and colleagues (2018) asked participants to retrieve either the stimulus color or the stimulus object in separate retrieval trials. Irrespective of the type of retrieval trial, encoding patterns were reliably reinstated in the occipitotemporal cortex. However, the type of trial could be decoded from patterns in the lateral parietal cortex, which represented the stimulus color during color trials and the stimulus object during object trials, thus flexibly adapting to the current retrieval goal. Therefore, manipulating task affordances during retrieval (e.g., attention, retrieval goal) may recruit additional regions in which distinctive neural representations are crucial to completing the task at hand. These additional regions may also be susceptible to age-related declines in neural distinctiveness, thus increasing the extent of observed age differences. However, more work is needed to investigate this hypothesis.

Above, we propose that retrieval-related age differences in neural specificity that are not attributable to reinstatement may be associated with a deficit in perceptual processing of stimuli during recognition in older adults. Active perceptual input during the old/new recognition task utilized in this study makes it difficult to definitively disentangle whether neural dedifferentiation during retrieval is related to poorly reinstated encoding patterns or poor sensory representations. While age effects of retrieval-related specificity partially overlapped with reinstatement age effects, age differences with regard to retrieval specificity were more widespread, potentially resulting from poor perceptual discriminability during recognition. Alterations in the neural representations of visual input in older adults may be a downstream consequence of poor sensory function (Lindenberger & Baltes, 1994; Schneider & Pichora-Fuller, 2000; Lindenberger et al., 2001; KZH Li & Lindenberger, 2002) and/or age differences in eye movements (Wynn et al., 2018; Wynn et al., 2019; Wynn et al., 2021). Age-related declines in both visual and auditory acuity have been associated with declines in cognitive performance (for review, see KZH Li & Lindenberger, 2002). Deterioration of sensory abilities likely leads to noisy and less specific neural representations in older adults (Schneider & Pichora-Fuller, 2000), which we may be observing in the decline in specificity during recognition. Similarly, age differences in eye movements may also be reflected in neural representations in visual areas. Wynn and colleagues (2021) observed that older adults had less distinctive gaze patterns across distinct images compared with younger adults, and when viewing the same image multiple times, older adults tended to gaze upon the same parts of the image with each presentation, while younger adults viewed new regions, thus updating and expanding their representations of the image. Both of these viewing behaviors in older adults were associated with poorer recognition memory performance (i.e., participants who had less distinctive gaze patterns tended to have worse recognition memory). In our view, age differences in gaze patterns may potentially feed forward into the specificity of neural representations in visual cortices. It would therefore be an interesting avenue to explore whether age differences in neural distinctiveness of visual representations are potentially mediated by age differences in eye movements.

Theories of cognitive aging (S-C Li & Lindenberger, 1999; S-C Li et al., 2001; for reviews, see S-C Li & Rieckmann, 2014; Koen & Rugg, 2019) hypothesize that neural dedifferentiation impairs memory performance. Although several studies have recently demonstrated a positive relationship between neural specificity during encoding and memory performance (Zheng et al., 2018; Koen et al, 2019; Srokova et al., 2020), retrieval-related specificity has not yet been associated with an objective measure of memory performance. We sought to close this gap by relating memory performance to our multivariate measure of specificity during retrieval. We did indeed identify a positive relationship between specificity and memory during retrieval. However, our findings suggest a far more prominent role of reactivation of categorical representations for successful mnemonic processing than specificity during retrieval. This aligns with several previous aging studies also reporting the beneficial impact of precise reinstatement on memory performance (St-Laurent et al., 2014; Abdulrahman et al., 2017; Bowman et al., 2019; Hill et al., 2021; but see Wang et al., 2016).

Further evidence pointing to the importance of reinstatement for successful memory comes from our trial-wise analyses showing that the strength of item-level reinstatement reliably predicted trial-by-trial memory outcomes. We additionally revealed a slight interaction in this relationship with age group, showing that item reinstatement was a predictor of memory success for younger adults, but not for older adults. This outcome resembles findings reported by Hill and colleagues (2021), who also showed an analogous age-moderated relationship between trial-wise category-level reinstatement and source memory performance for scene stimuli (but, see Trelle et al., 2020, for findings of a trial-wise relationship between memory success and item reinstatement in older adults). In contrast to both Trelle et al. (2020) and Hill et al. (2021), we did not find that retrieval-related hippocampal activity predicted memory success. This may be yet another manifestation of how retrieval task influences the results, since both of the aforementioned studies used a source memory retrieval task, which is more likely to recruit hippocampal resources to support memory retrieval through pattern completion (McClelland et al., 1995).

Age-related neural dedifferentiation has also been reported in terms of a decrease in the distinctiveness of item-specific reinstatement (Folville et al., 2020; Hill et al., 2021). Although we observed occipital regions demonstrating an effect of strong item-level specificity during reinstatement for houses, we found neither age differences in this effect nor an interindividual relationship with memory performance. The absence of age differences in item-level reinstatement is surprising and appears to contradict previous evidence for an age deficit at this representational level. However, while Hill and colleagues (2021) demonstrated an age-related decrease in item-level pattern similarity, their measure did not control for potential age deficits at the category level. Therefore, the observed age differences at the item level may not have been reflected more than a general categorical deficit. Nevertheless, our finding proves difficult to interpret in the context of the current literature — more studies will be needed to understand how age differences in neural distinctiveness vary across different representational levels.

Finally, age differences in neural distinctiveness were found in this study for houses, but not for faces. Age-related neural dedifferentiation has frequently been documented for “place” stimuli, such as houses and scenes (Voss et al., 2008; Carp et al., 2011; Zheng et al., 2018; Koen et al., 2019; Srokova et al., 2020, Hill et al., 2021), with one exception (Berron et al., 2018). However, evidence for dedifferentiation for faces is more mixed, with some studies reporting age differences (D.C. Park et al., 2004; J. Park et al., 2012; Voss et al., 2008) and some reporting absent age effects (Payer et al., 2006; Srokova et al., 2020; Hill et al., 2021). Our findings mirror those of Hill and colleagues (2021), who reported age differences in category-level reinstatement and category specificity during retrieval only for scene stimuli and not for face stimuli. The present discrepancy between face and house stimuli may be attributed to a couple of factors. First, it has been argued that age differences may be less salient for faces due to the lifetime experience hypothesis (Koen et al., 2019; Srokova et al., 2020), which posits that perceptual schemas may develop across the life course as a result of substantial exposure to particular categories, possibly allowing for more rapid integration of novel exemplars into these schemas. Thus, absent age differences for faces may reflect that the face schema is already developed by young adulthood. In addition, categorical differences may stem from lower order feature-level differences. For example, houses have been shown to be more feature-rich than faces, which drives brain signal variability (Garrett et al., 2020) and may also impact measures of neural representations. Therefore, the neural dedifferentiation field could benefit from studies incorporating additional stimulus categories that vary at the feature level in order to disentangle whether these observed disparities between face and house stimuli are actually category-specific effects or rather related to lower-order stimulus features.

Together, our findings reveal evidence for age-related neural dedifferentiation during memory retrieval. Importantly, these age deficits cannot be fully attributed to imprecise reinstatement of encoding-related category representations and thus may also reflect impaired perceptual processing in older adults. Furthermore, while both retrieval- and reinstatement-related category specificity were positively associated with memory performance across participants, reinstatement specificity correlated with memory in substantially more neural regions than retrieval specificity. We propose that these outcomes highlight the significance of precise cortical reinstatement for the support of memory retrieval. We also suggest several avenues for future research, including investigation of how the retrieval task might play a role in manifestations of age-related neural dedifferentiation.

## Acknowledgments

We thank Verena R. Sommer and Attila Keresztes for programming and running the study, all student assistants who helped with data collection, Gabriele Faust and members of the LIME and RHYME projects for helpful feedback, Julia Delius for editorial assistance, and all study participants for their time.

